# Little variation in the morphology of the atria across 13 orders of birds

**DOI:** 10.1101/397034

**Authors:** Jelle G. H. Kroneman, Jaeike W. Faber, Claudia F. Wolschrijn, Vincent M. Christoffels, Bjarke Jensen

## Abstract

Mammals and birds acquired high performance hearts and endothermy during their independent evolution from amniotes with many reptile characters. A literature review shows that the variation in atrial morphology is greater in mammals than in ectothermic reptiles. We therefore hypothesized that the transition from ectothermy to endothermy associated with greater variation in cardiac structure. We tested the hypothesis in birds, by assessing the variation in 15 characters in hearts from 13 orders of birds. Hearts were assessed by gross morphology and histology, and we focused on the atria as they have multiple features that lend themselves to quantification. We found bird hearts to have multiple features in common with ectothermic reptiles (synapomorphies), for instance the presence of three sinus horns. Convergent features were shared with crocodylians and mammals, such as the cranial offset of the left atrioventricular junction. Other convergent features like the compact organization of the atrial walls were shared with mammals only. Sinus myocardium expressing Isl1 was node-like (Mallard), thickened (chicken), or anatomically indistinct from surrounding myocardium (Lesser redpoll). Some features were distinctively avian (apomorphies), including the presence of a left atrial antechamber, and the ventral merger of the left and right atrium, which was found in parrots and passerine birds. Most features, however, exhibited little variation. For instance, there were always three systemic veins and two pulmonary veins, whereas among mammals there are 2-3 and 1-7, respectively. Our findings suggest that the transition to high cardiac performance does not necessarily lead to greater variation in cardiac structure.

## 1 INTRODUCTION

Mammals and birds evolved independently from reptile-like ancestors as two vertebrate classes that are characterized by high metabolic rates and endothermy (Warren, 2008; Green., et al., 2014; Tattersall, 2016). When the hearts of mammals and reptiles are compared, it is evident that mammalian hearts are remodeled by incorporation of systemic and pulmonary vein myocardium to the atria (Jensen, Boukens, Wang, Moorman, Christoffels, 2014a; Carmona, Ariza, Cañete, Muñoz-Chápuli, 2018). The number of venous orifices to the left atrium can vary between 1 (dugongs) and 7 (armadillos), and the myocardial sleeve of the veins may be extensive (mouse) or all but gone (harbour porpoise) (Rowlatt, 1990; Mommersteeg et al.,2007). The number of orifices and the extent of myocardium even varies within the species (Nathan, Eliakim, 1966; Calkins et al., 2007, Rowlatt, 1990). In human, for example, the left atrium typically receives four pulmonary veins, but three and five veins are also frequently observed, two and six can occur but are rare (Mansour et al., 2004). In contrast, in reptiles the pulmonary circulation connects to the left atrium by a single orifice (Jensen, Moorman, Wang, 2014b). These examples could suggest that the transition from ectothermy to endothermy, and the associated rise in cardiac pumping (Crossley et al, 2016), initiated greater variation in the building plan of the heart. We therefore hypothesized that birds will have hearts that exhibit similar degrees of variation as mammals.

The extensive studies of the chicken heart, its development in particular, sharply contrasts the comparatively meagre number of anatomical studies on hearts of other birds (Hamburger, Hamilton, 1951; Van Mierop, 1967; Poelmann, Mikawa, Gittenberger-De Groot, 1998; Sedmera, Pexieder, Vuillemin, Thompson, Anderson, 2000; Lincoln, Alfieri, Yutzey, 2004; van den Berg et al., 2009; Bressan, Lui, Louie, Mikawa, 2016). Literature on the venous-atrial region in avian species is limited (Jensen et al., 2014a). In chicken, there is a myocardial sleeve surrounding the systemic and pulmonary veins that are within the pericardial cavity (Endo, Kurohmaru, Nishida, Hayashi, 1992; van den Hoff, Kruithof, Moorman, Markwald, Wessels, 2001), but similar studies have, to the best of our knowledge, not been extended to other avian species. Older studies made gross anatomical comparisons of multiple species (Gasch, 1888; Benninghoff, 1933) but a general paucity of images and quantifications makes it difficult to verify these findings, let alone to make comparisons to other clades of vertebrates. For instance, the walls of the atria are described as thin with thick bundles of muscle forming muscular arches (Whittow, 1999; Sedmera *et al*., 2000), but how this setting compares to other vertebrates remains difficult to assess. We therefore undertook a study to assess the variation in the morphology of the avian heart. We focused on the atria and the base of the ventricle, to assess the 15 features schematized in Figure 1 across 13 orders of birds.

**Figure 1.**
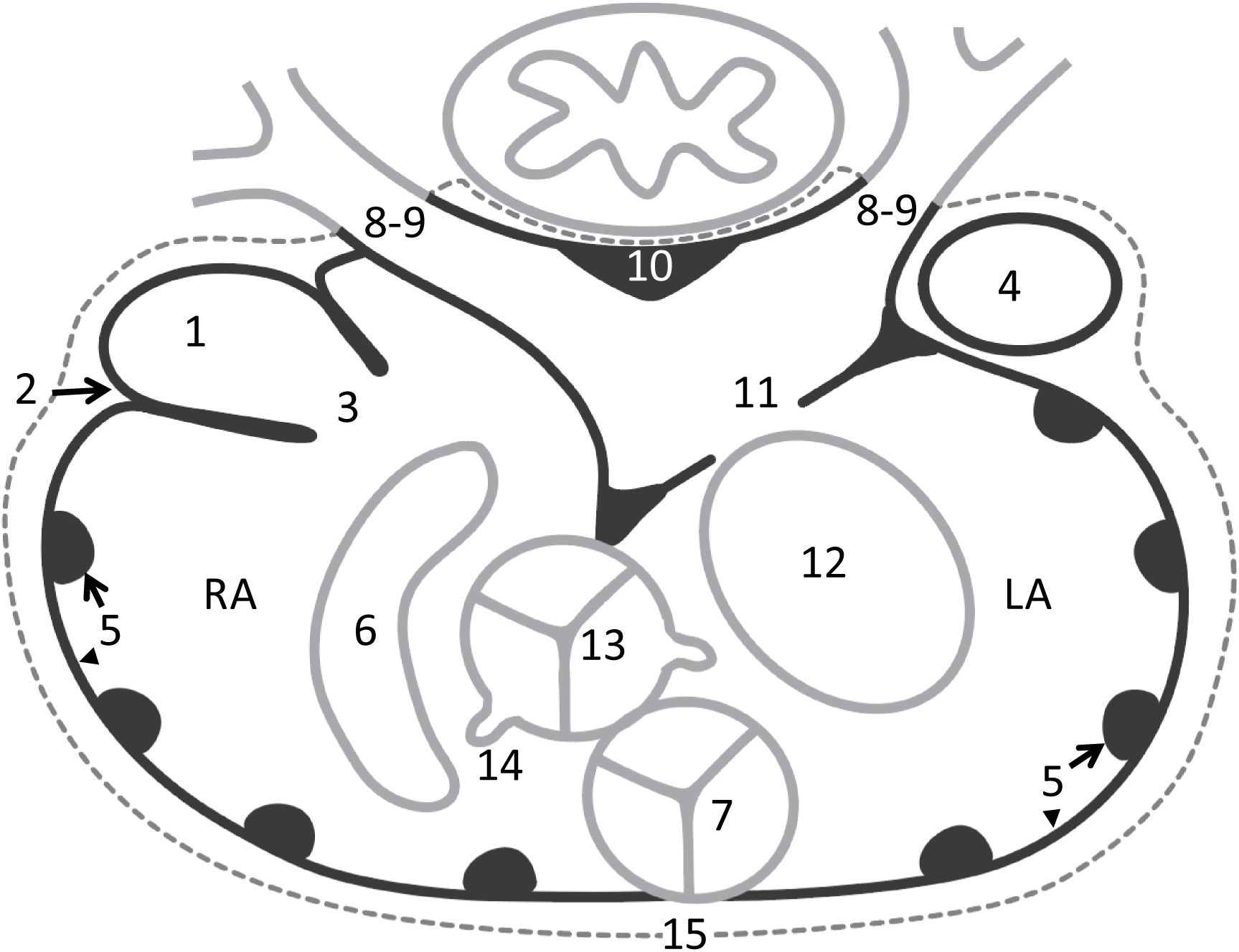
Scheme of the analyzed structures of the avian hearts. 1) myocardium in the sinus venosus, 2) the sinuatrial node, 3) sinuatrial valve and leaflets, 4) left sinus horn, 5) pectinate muscles of the atrial wall, 6) position of right atrioventricular junction and muscularity of the valve, 7) position of the pulmonary artery and valve leaflets, 8) number of pulmonary veins, 9) the extent of pulmonary venous myocardium relative to the pericardium (dashed line), 10) dorsal ridge of the antechamber of the left atrium, 11) muscular shelf in the roof of the left atrium, 12) position of left atrioventricular junction and muscularity of the valve, 13) position of the aorta and valve leaflets, 14) position of the orifices of the coronary arteries, 15) presence of ventral merger of the atrial walls.

## 2 MATERIALS AND METHODS

### 2.1 Isolated hearts

One heart had been isolated from one specimen of the Ostrich (*Struthio camelus*, Struthioniformes) taken from a previously published model (Jensen et al., 2013). The heart of the Mallard (*Anas platyrhynchos*, Anseriformes) was taken from a culled animal. The heart of the adult Chicken (*Gallus gallus*, Galliformes) came from an animal killed for human consumption, and the heart from a developing chicken (Hamburger-Hamilton stage 44) was isolated from an in-house incubated embryonated egg (Amsterdam UMC). The heart of the Lesser redpoll (*Acanthis cabaret*, Passeriformes) was isolated from an animal that was found dead on a bike path (BJ). The heart of the Collared dove (*Streptopelia decaocto,* Columbiformes), Common swift (*Apus apus*, Apodiformes), Eurasian coot (Fulica Atra, Gruiformes), Common snipe (*Gallinago gallinago*, Charadriiformes), Sparrowhawk (*Accipiter nisus*, Accipitriformes), Barn owl (*Tyto alba,* Strigiformes), Green woodpecker (*Picus viridis,* Piciformes), Common kestrel (*Falco tinnunculus*, Falconiformes), Budgerigar (*Melopsittacus undulatus*, Psittaciformes), Barn swallow (*Hirundo rustica,* Passeriformes), and Hawfinch (*Coccothraustes coccothraustes*, Passeriformes) came from the Department of Pathobiology (Utrecht University). The age is not known for any specimen, except for the HH44 chicken, but all animals were deemed to be of approximate adult size.

### 2.2 Tissue preservation

The adult Mallard, Chicken HH42, and Lesser redpoll were fixed in 4% PFA for 24 hours and then stored in 70% ethanol. The other samples came from hearts used in anatomy classes at the Department of Pathobiology: these had initially been frozen in -18 °C, thawed in water, fixed in 4% formaldehyde solution, and afterwards stored in 1% formaldehyde solution. Before use in class, the samples were rinsed for 48 hours with running tap water. This process had occurred multiple times with the exact amount unknown to us.

### 2.3 Histology and immunohistochemistry

The hearts from Barn swallow, Green woodpecker, Collared dove, Chicken, Budgerigar, Common snipe, Common swift, Grey heron, Barn owl, Common kestrel and Eurasian coot were embedded in paraplast and cut in 10 µm transverse sections, apart from the Collared dove which was cut in 4-chamber view. The principal staining was picrosirius red (muscle is stained orange, collagen red with 2 min differentiation in 0.01 M HCL). The atrial region of each heart may yield several hundreds of sections and we selected, at a fixed distance (dependent on the size of the heart), approximately 20 sections to represent the entire atrial region from below the atrioventricular junction to the roof of the atria. Using immunohistochemistry, we detected myocardium with cTnI mouse antibodies (Invitrogen dilution 1:200) or cTnI rabbit antibodies (Hytest 1:200) visualized by a fluorescently labelled secondary donkey-anti-mouse antibody (Invitrogen, dilution 1:200) or donkey-anti-rabbit antibody (Invitrogen, dilution 1:200) respectively coupled to Alexa 488. Arterial musculature was detected with a mouse antibody to smooth muscle actin (Millipore, dilution 1:250) visualized by a fluorescently labeled secondary donkey anti-mouse antibody (Invitrogen, dilution 1:200) coupled to Alexa 568. For the identification of the sinuatrial node, Isl1 goat antibodies (Neuromics, dilution 1:200) visualized by a fluorescently labelled secondary donkey-anti-goat antibody coupled to Alexa 647 (Invitrogen, dilution 1:200) were used. Nuclei were stained with sytox orange (Invitrogen, dilution 1:500). *In situ* hybridization was performed as described previously (Jensen *et al*. 2017), with probes against cardiac troponin I (*cTnI*) and bone morphogenetic protein 2 (*Bmp2)* (Somi, Buffing, Moorman, van den Hoff, 2004).

### 2.4 Imaging

Imaging of the picrosirius red stained slides was done with a Leica DM5000 light microscope. For the larger sections, we merged multiple photos using the ‘Photomerge’ function in Adobe Photoshop CS6, version 13.0.1. Several of the hearts contained large quantities of blood that stained in a color similar to that of the muscular walls. In many instances, to ensure clarity, these areas were masked with white to make the cardiac tissue stand out. If this was done, it is stated in the figure legend. For immunohistochemistry, slides were viewed and photographed with a Leica DM6000B fluorescent microscope. For the Barn swallow, Green woodpecker, and the Mallard, all analyzed structures were annotated in Amira software (version 6.0.0, FEI, SAS). To get volume estimates of each structure, the ‘Materials Statistics’ tool in Amira was used (Supplementary Figure 1). These three specimens were chosen because of the quality of the section series and because they span two orders of magnitude in heart and body size (approximately 10, 100, and 1000g).

### 2.5 Analyzed structures

Figure 1 shows the 15 structures that were analyzed: (1) Presence of myocardium in the sinus venosus, (2) location of the sinus node, (3) the sinuatrial valve and the number of leaflets it contains, (4) existence of a left sinus horn, (5) presence and size of pectinate muscles in the atrial wall, (6) anatomy and position of the right atrioventricular valve apparatus, (7) anatomy of the pulmonary arterial valve, (8) number of pulmonary veins entering the left atrium, (9) extent of pulmonary venous myocardium, (10) the dorsal myocardial ridge in the antechamber of the left atrium, (11) the muscular shelf in the left atrial roof, (12) anatomy and position of the left atrioventricular valve apparatus, (13) anatomy of the aortic valve, (14) origins or the coronary arteries, (15) presence of ventral merger of the left and right atrial walls ventral to the pulmonary trunk.

All hearts were gross morphologically inspected and were assessed on the basis of sections. Due to damage, not all structures could be assessed in all hearts. Supplementary Table 1 lists for each heart which structures were investigated and how.

**Table 1.**
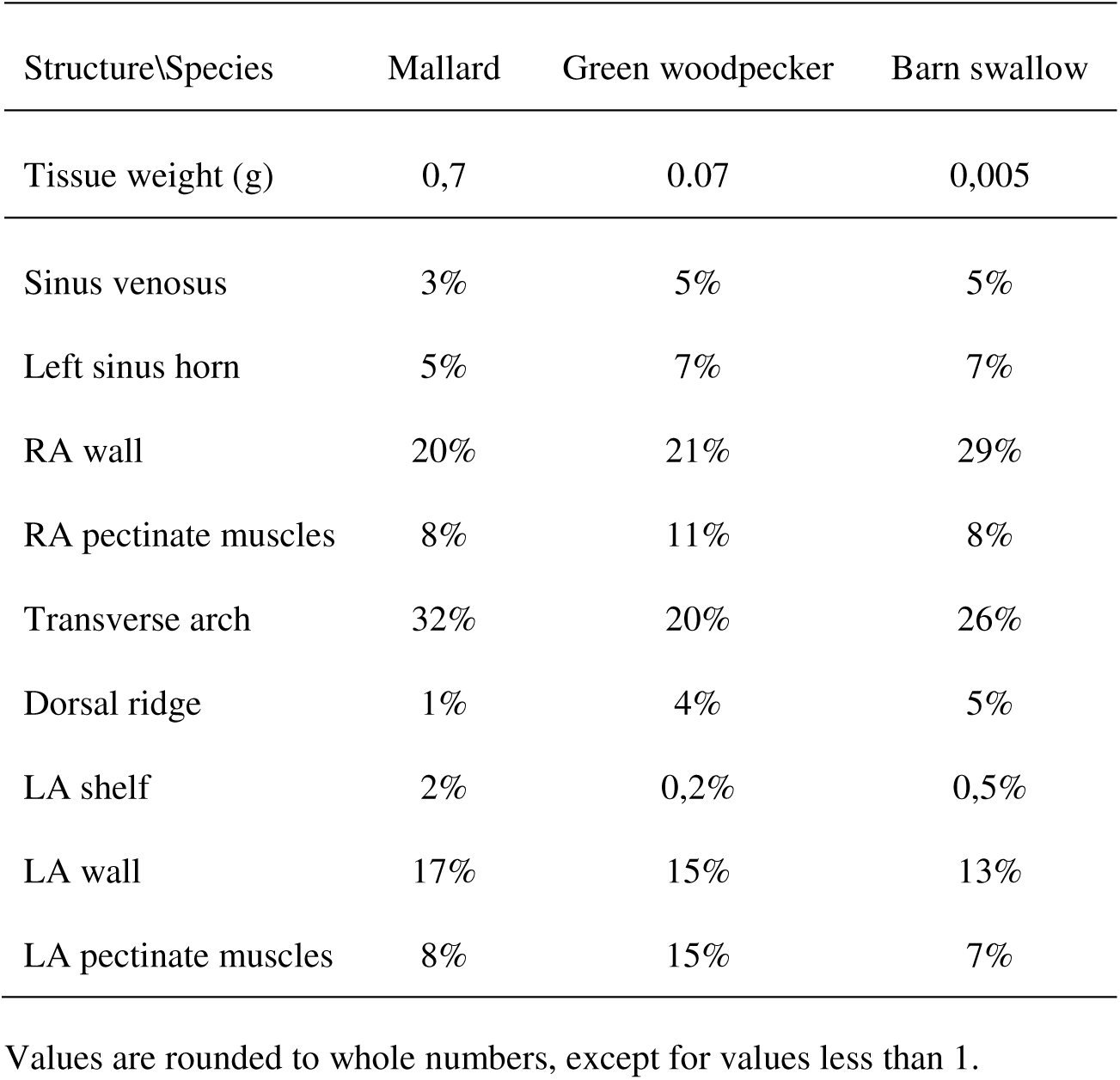
Proportions of atrial structures, estimated from areas in Amira.

| Structure\Species | Mallard | Green woodpecker | Barn swallow |
| --- | --- | --- | --- |
| Tissue weight (g) | 0,7 | 0.07 | 0,005 |
| Sinus venosus | 3% | 5% | 5% |
| Left sinus horn | 5% | 7% | 7% |
| RA wall | 20% | 21% | 29% |
| RA pectinate muscles | 8% | 11% | 8% |
| Transverse arch | 32% | 20% | 26% |
| Dorsal ridge | 1% | 4% | 5% |
| LA shelf | 2% | 0,2% | 0,5% |
| LA wall | 17% | 15% | 13% |
| LA pectinate muscles | 8% | 15% | 7% |

Values are rounded to whole numbers, except for values less than 1.

## 3 RESULTS

In all investigated hearts, the heart was dorsal to the sternum. The cardiac long-axis was parallel to the sternum and the sternal carina which ran parallel to the spine. The apex of the cardiac ventricles was the caudal most point of the heart. All major arteries projected cranial from the ventricular base, and the atria were cranial to the ventricles. Tissue close to the sternum was considered ventral, the tissue closest to the spine was considered dorsal.

### 3.1. Gross morphology

The structures of the bird atria are described in the sequence in which the blood courses through the heart, starting at the right side. Three large systemic veins enter the pericardial cavity, the right cranial vena cava, the left cranial vena cava, and the caudal vena cava. Their walls contain myocardium and are, therefore, considered to be derivatives of the sinus venosus, the chamber upstream of the right atrium in ectotherms. This is why, the three vessels will be further referred to as the right sinus horn, left sinus horn, and caudal sinus horn. The opening of the sinus venosus into the right atrium always has a myocardial valve, comprising a left and a right leaflet. The luminal side of the wall of the right atrium has multiple pectinate muscles of varying sizes. The pectinate muscles converge cranially in a large muscular arch that spans the roof of both atria from right to left, the so-called transverse arch (it constitutes approximately a quarter of atrial muscle, see below). At the bottom of the right atrium, the entrance to the right ventricle is guarded by the large muscular flap valve, which, on the ventricular side, is made up of thick ventricular wall, and on the atrial side of much thinner atrial muscle, with a thin layer connective tissue between the atrial and ventricular myocardium.

The left atrium receives two pulmonary veins in an antechamber before opening into the body of the left atrium. This antechamber consists of the myocardial walls of the pulmonary veins which have a dorsal ridge of myocardium between them which is oriented parallel to the esophagus. A muscular shelf in the roof of the atrial cavity partly separates this antechamber from the body of the left atrium. The walls of the antechamber are smooth, as opposed to the walls of the body of the left and right atrium, which are both trabeculated. A single atrial septum, without a *fossa ovalis*, separates the left and right atrium. This septum is continuous with the muscular shelf. The right atrium appears more voluminous than the left atrium, in part due the somewhat left-ward position of the atrial septum in combination with a greater caudo-cranial height of the right atrial cavity. The entrance to the left ventricle is guarded by the mitral valve consisting of thin fibro-membranous leaflets. From histology, we could not assert with certainty the number of leaflets. The outflow tract of both ventricles, the aorta and pulmonary artery, are partly embraced by the atria and are situated ventrally, the pulmonary artery being the most ventral.

### 3.2 The sinus venosus

Myocardium was found in the left sinus horn, right sinus horn, and caudal sinus horn (Figure 2) (myocardium was principally assessed on picrosirius red stained sections, complemented with immunohistochemistry in a few sections per section series). The amount of myocardium varied between sinus horns. In all birds in which histology was performed (see Supplementary Table 1), the myocardium of all three sinus horns stops in proximity of the pericardium. In the birds analyzed with Amira, the volume of the sinus venosus was compared to the volume of the right atrium (right atrial wall and right pectinate muscles) and was found to be approximately 12% (Table 1, Supplementary Figure 1).

**Figure 2.**
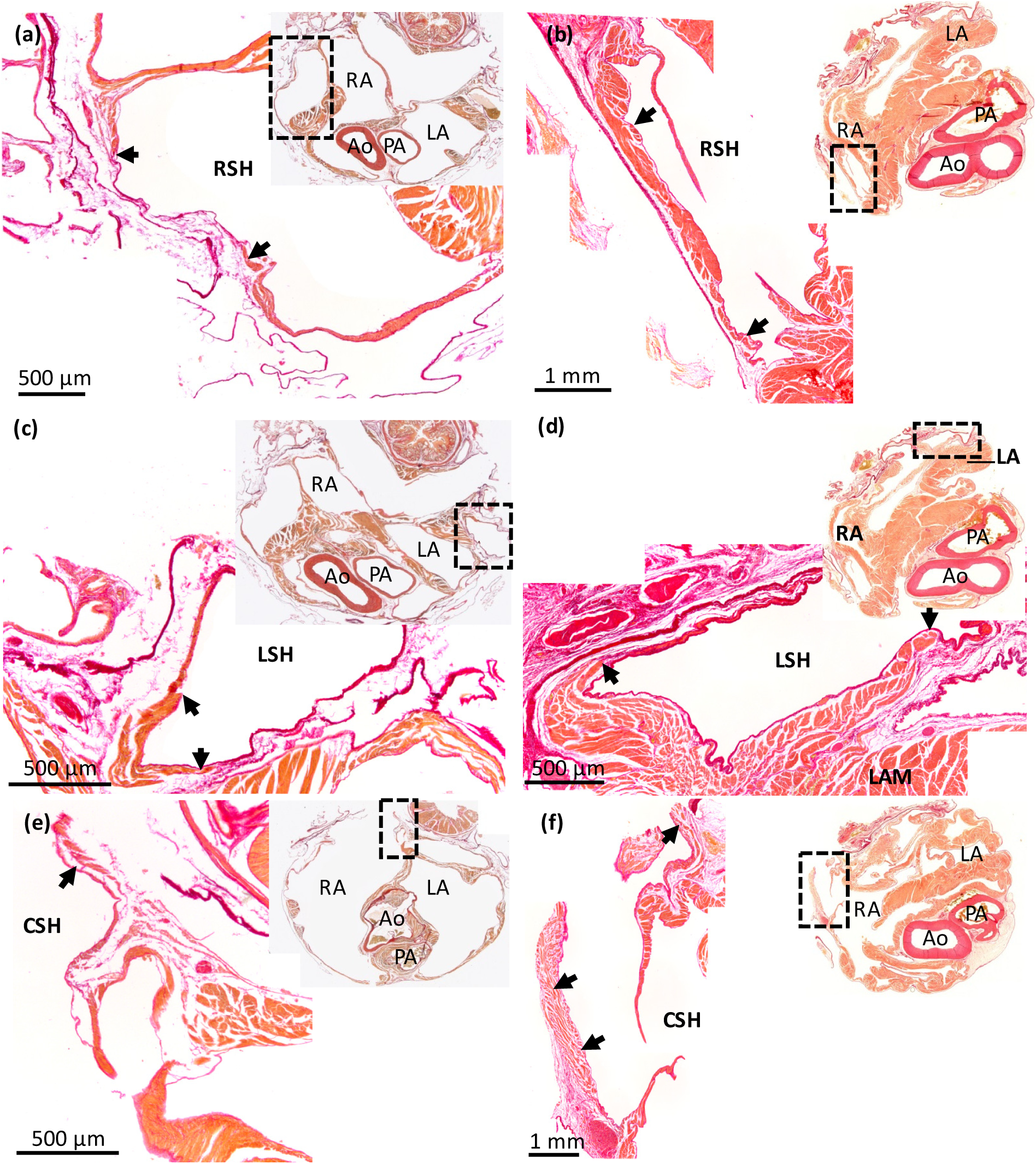
Myocardium in the sinus venosus of the Budgerigar (a,c,e) and the Mallard (b,d,f). Picrosirius red stained 10µm histological sections. The right sinus horn (RSH) (a,b), left sinus horn (LSH) (c,d), and caudal sinus horn (CSH) (e,f) contain myocardium (black arrows). The myocardial wall can be quite thick proximal to the atria, but tapers off distally. At the pericardial border, the vessel wall may be without myocardium. Ao, aorta; LA, left atrium; LAM, left atrial muscle; PA, pulmonary artery; RA = right atrium. In the images from the Budgerigar, blood has been painted over with white for clarity.

A left sinus horn was present in all bird hearts. Proximal to its entry to the right atrium, its myocardial sleeve was thick and fully surrounded the lumen (Figure 2). As the left sinus horn opened into the right atrium some muscle protruded into the lumen as a leaflet (Figure 3). Such a leaflet was observed in all birds apart from the Common kestrel where the leaflet was thin and membranous. Compared to the total volume of myocardium in the sinus venosus, the myocardium of the left sinus horn comprised approximately 60% (Table 1).

**Figure 3.**
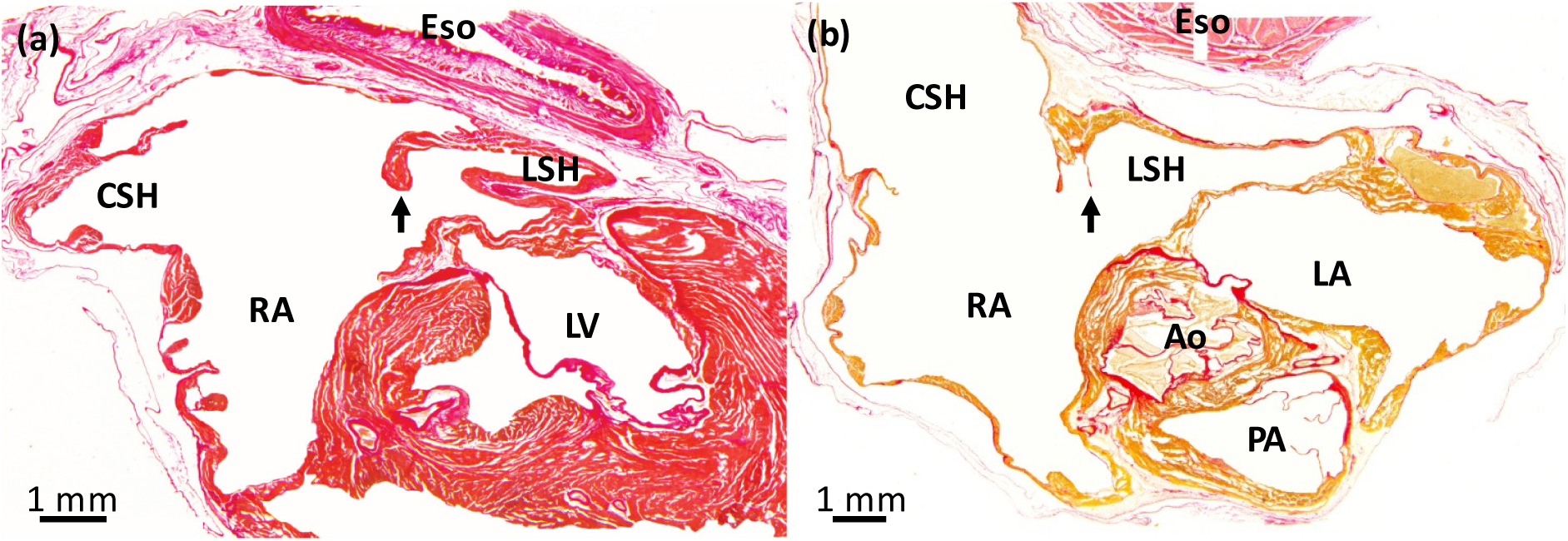
Leaflet of the valve of the left sinus horn in the Green woodpecker (a) and the Common kestrel (b). Birds generally have a prominent valve (black arrow) guarding the orifice of the left sinus horn (LSH), here exemplified by the Green woodpecker. It was found on sections representing 0.7mm out of the total height of the atria of 6.8mm. In the Common kestrel, the valve was very thin membranous leaflet and it was found on sections representing 0.7mm out of the total height of the atria of 6.6mm. Ao, aorta; CSH, caudal sinus horn; Eso, esophagus; LA, left atrium; LV, left ventricle; LSH, left sinus horn; PA, pulmonary artery; RA, right atrium. In the images, blood has been painted over with white for clarity.

The entrance of the right sinus horn into the right atrium is separate from that of the caudal sinus horn and left sinus horn in most birds. Only in the Green woodpecker, the right sinus horn is part of a larger sinus venosus. The orifice of the right sinus horn is guarded by the sinuatrial valve, the left leaflet of which was most developed in this region of the right atrium. Cranially, the left leaflet has a thick muscular margin that merges with the transverse arch in the Common swift, Eurasian coot, Green woodpecker, and Budgerigar (Supplementary Figure 2). Caudally, the right leaflet is more developed in the Mallard, Common kestrel and Barn swallow, whereas the left leaflet appears to be more developed in the Common swift, Eurasian coot and Budgerigar. The position of the orifice of the right sinus horn and caudal sinus venosus varied between species. We measured these positions relative to a dorsal-ventral axis that was established from the position of esophagus, atrial septum, aorta and pulmonary vein (Supplementary Figure 3).

### 3.3 Sinuatrial node in bird hearts

Detection of markers of the sinuatrial node was successful in adult Mallard, Chicken HH42, and Lesser redpoll, the best-preserved specimens (Figure 4). In the Mallard, Isl1 detection was confined to a small oval-shaped structure at the base of the right leaflet of the sinuatrial valve (Figure 4a-c, Supplementary Figure 4). At its most expansive, the sinuatrial node cross-section was approximately 700 µm wide and 900 µm long, detectable on eight sections each 300 µm apart, giving a total volume of approximately 1,5 mm^3^. A large coronary artery, identified by the expression of smooth muscle actin in the arterial wall, was found within the Isl1 positive domain (Figure 4b). This domain was relatively rich in collagen compared to the surrounding myocardium of the sinus venosus and right atrium (Figure 4a). In Chicken, the sinuatrial node is anatomically less distinct than in the Mallard. The base of the right leaflet of the sinuatrial valve has less expression of *cTnI* than the surrounding muscle (Figure 4d-e), and it was the only muscle that expressed *Bmp2* (Figure 4f) and Isl1 (Figure 4g). In the Lesser redpoll, there was no anatomically identifiable node (Figure 4h). The base of the right leaflet of the sinuatrial valve is thicker than the surrounding walls, and this region expresses Isl1 (Figure 4i-j) and harbors a large coronary artery (Figure 4i). In most specimens, however, Isl1 could not be detected. On the picrosirius red stained sections with reasonably good to good tissue presentation, a Mallard-like sinus node was found in the budgerigar only (Supplementary Table 1). In the swift, woodpecker, and kestrel, there was no mallard-like sinus node, but rather thickened myocardium on the sinus-side of the right leaflet of the sinuatrial valve which resembled the Isl1 positive domain of the Lesser redpoll (Supplementary Table 1). These three species are phylogenetically found in between the mallard and budgerigar, and there seems to be no obvious phylogenetic trend concerning the anatomy of the putative sinus node.

**Figure 4.**
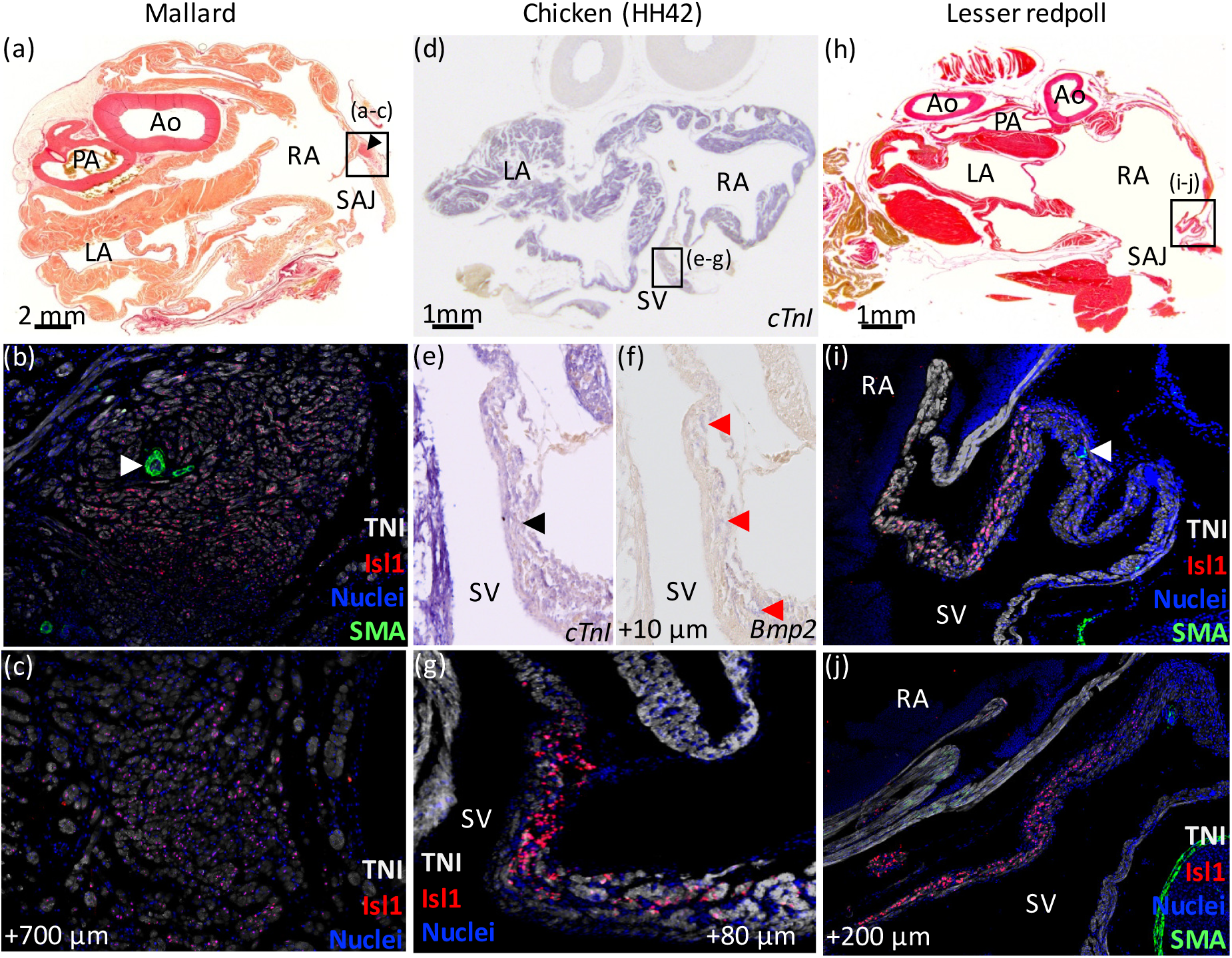
Sinuatrial node in Mallard (a-c), Chicken HH42 (d-g), and Lesser redpoll (h-j). (a-c) In the Mallard, there was a nodal structure at the base of the right leaflet of the sinuatrial valve (black arrowhead in (a)) which expressed Isl1 (b-c) and which had a large coronary artery (white arrowhead in (b)). (d-g) In the Chicken HH42, the sinus venosus (SV) expressed the myocardial marker *cTnI* and this expression was relatively weak at the base of the right leaflet of the sinuatrial valve (black arrowhead in (e)). (f-g) The base of the right leaflet of the sinuatrial valve expressed *Bmp2* (red arrowheads in (f)) and Isl1 (g). (h-j) In the Lesser redpoll, Isl1 was expressed in the base of the right leaflet of the sinuatrial valve. There was no nodal structure but the Isl1 expressing wall was thicker than the surrounding walls and contained a large coronary artery (white arrowhead in (i)). Ao, aorta; LA, left atrium; PA, pulmonary artery; RA, right atrium; SAJ, sinuatrial junction.

### 3.4 The right atrium

In all bird hearts, extensive pectinate muscles were found in the right atrial wall. The smallest birds, Barn swallow and Lesser redpoll, have the fewest pectinate muscles, 4 and 5 respectively, whereas the much larger Mallard had approximately 18 across the atria. Most pectinate muscles were nodular in cross section and much thicker than the atrial wall (Figure 5). In Budgerigar, a large trabecula was 0.56 mm while the atrial wall next to it was 0.021 mm; this means the trabecula was 27 times thicker than the atrial wall next to it. In Mallard, a large trabecula was 1.79 mm while the atrial wall next to it was 0.22 mm so the trabecula was 8 times thicker than the atrial wall next to it (Figure 5). The pectinate muscles connected together in a large muscular transverse arch in the roof of both atria. In the birds analyzed with Amira (Table 1), half of the atrial muscle was trabeculated (transverse arch, left atrial pectinate muscles, right atrial pectinate muscles and dorsal ridge). The atrial septum was thin and we never observed a shallow depression akin to the foramen ovale of eutherian mammals.

**Figure 5.**
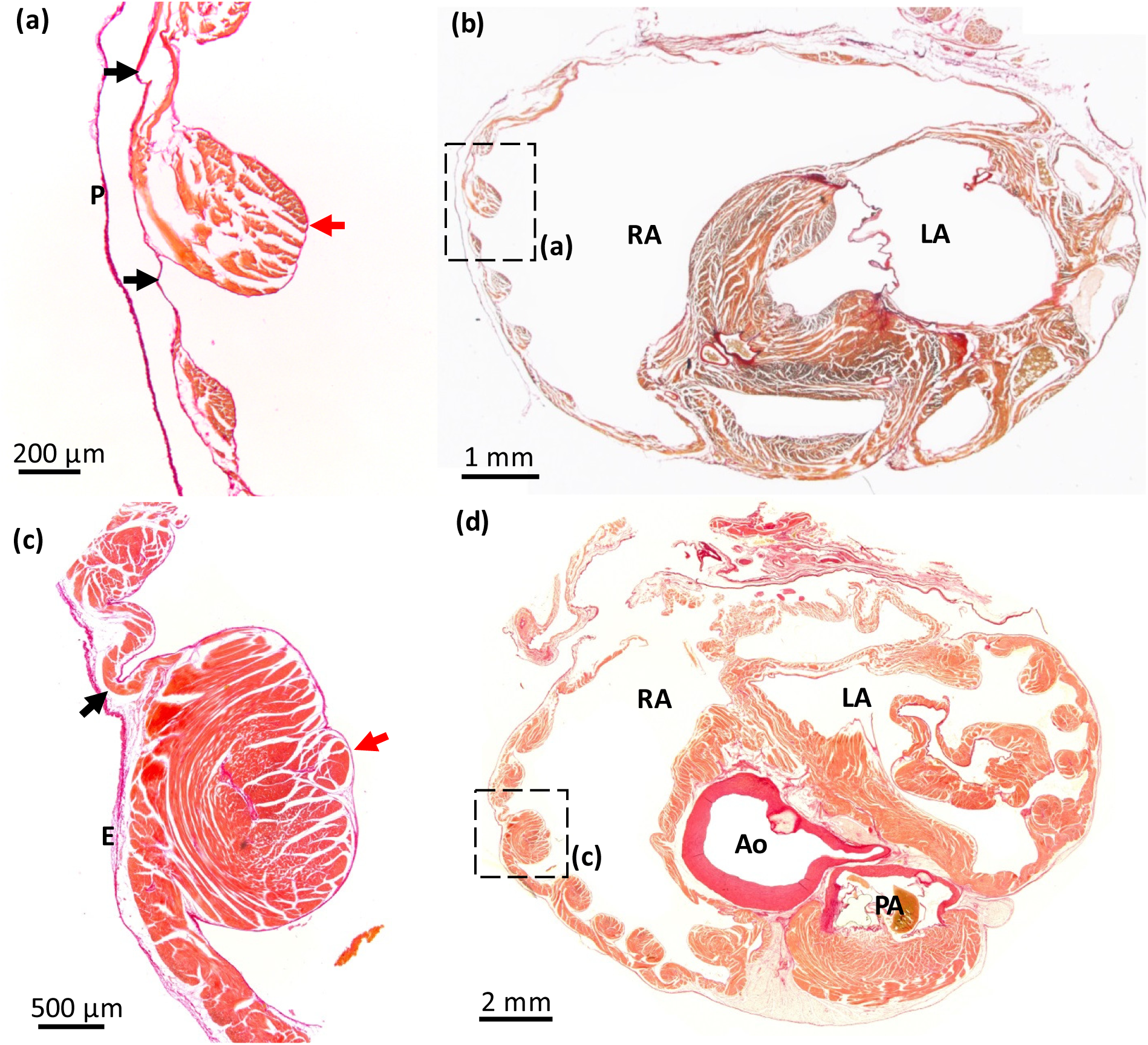
Atrial pectinate muscles in the Barn swallow and the Mallard. Picrosirius red stained section of Budgerigar (a,b) and Mallard (c,d). The position of the pectinate muscles imaged in (a,c) are indicated by black squares in the overviews (b,d). Red arrows indicate pectinate muscles, black arrows point to the atrial wall in between the pectinate muscles. Note the several-fold greater thickness of the pectinate muscles to the atrial wall. Ao, aorta; E, epicardium; LA, left atrium; P, pericardium; PA, pulmonary artery; RA, right atrium. In the images from the Budgerigar, blood has been painted over with white for clarity.

### 3.5 Position of the atrioventricular orifices

In transverse sections, the atrioventricular and arterial orifices were seen to be nestled together (Figure 6). The aortic valve was almost in the center of the ventricular base. Relative to this position, the right atrioventricular junction was dorsal and right, the left atrioventricular junction was dorsal left, and the pulmonary arterial valve was located ventrally (Figure 6). The right atrioventricular junction was configured as a C, with the medial margin provided by the interventricular septum and the parietal margin provided by the large muscular flap valve. Ventrally, the flap valve merged with the septal surface. The left atrioventricular junction was always rounded and guarded by a valve of connective tissue (Figure 6). When we inspected the histological sections from cranial to caudal, the leaflets of the left atrioventricular junction always appeared before the flap valve of the right atrioventricular junction (Supplementary Table 2), showing there was always an caudo-cranially offset between the two atrioventricular junctions (Supplementary Figure 5). The left atrioventricular junction appeared 1.65mm before the right atrioventricular junction averaged over 9 species. Regarding this feature, the Eurasian coot had the largest offset at 4,4mm and the Common snipe had the smallest with 0.3mm.

**Figure 6.**
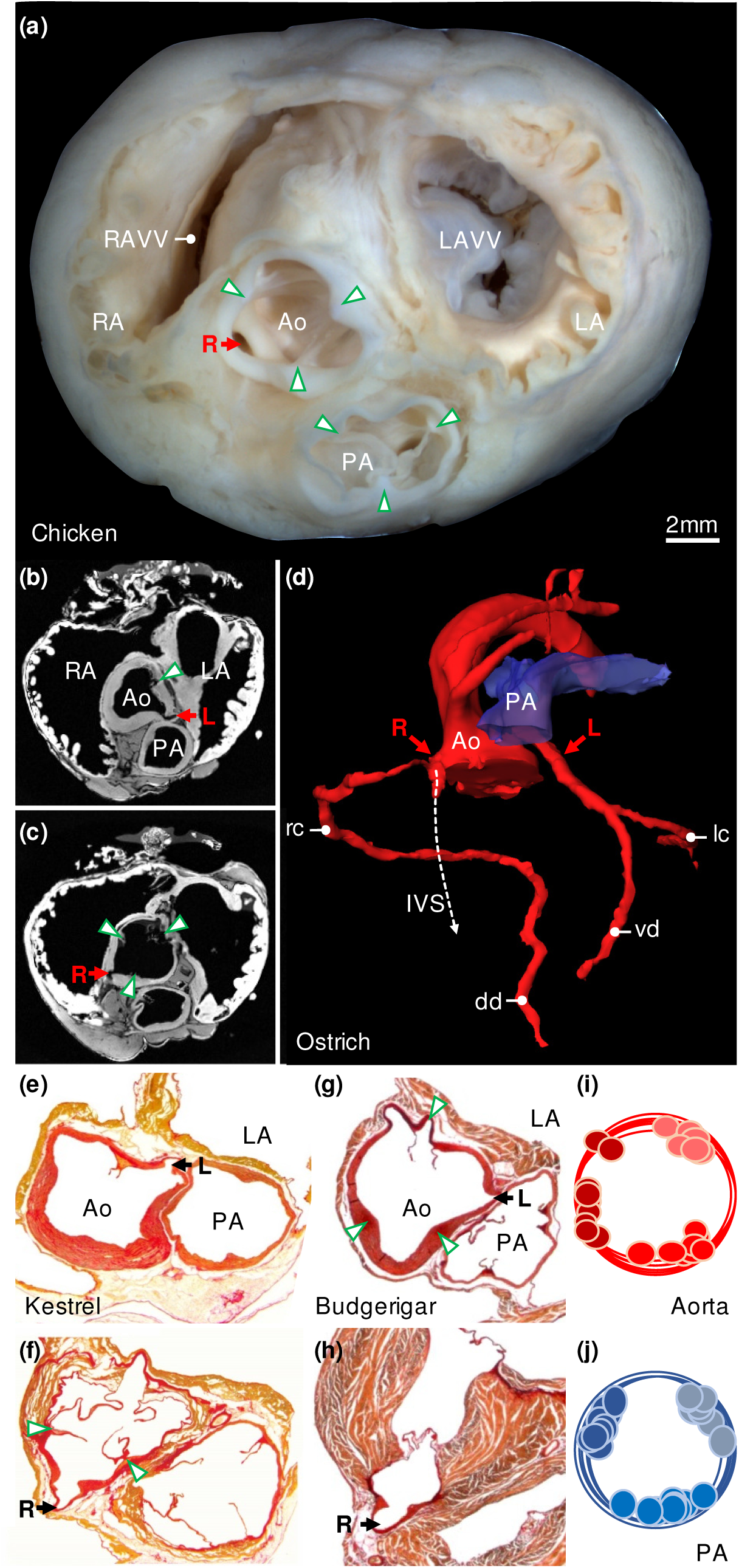
Major arteries. (a) Gross morphology of the ventricular base of the with indication of the valve commissures (green arrowheads) and the right coronary artery (R). (b-d) In the Ostrich, the left coronary artery (L) originates from the sinus of the ventral-left leaflet of the aortic valve (b) and the right coronary artery (R) originates from the sinus of the ventral-right leaflet of the aortic valve (c) (arrow heads indicate leaflet commissures). (d) The 3D reconstruction of the lumen of the major arteries shows the right coronary immediately splits into a branch that runs in the interventricular septum (IVS), next to the ventral merger of the right atrioventricular valve and the interventricular septum. The right coronary gives rise to the right circumflex artery (rc) leading to the dorsal descending artery (dd) in the interventricular sulcus. The left coronary splits into the ventral descending artery (vd) in the interventricular sulcus and the left circumflex artery (lc). The images of (b-d) are derived and modified from the Ostrich 3D model published previously (Jensen et al., 2013). In the Common kestrel (e-f) and Budgerigar (g-h), the origin of the coronary arteries from the aorta was like in the Ostrich, and the intraseptal branch of the right coronary artery was always found to be located next to the ventral merger of the right atrioventricular valve with the interventricular septum. The cartoons (i-j) show the distribution of the commissures of the valve leaflets between all sectioned birds. Ao, aorta; LA, left atrium; LAVV, left atrioventricular valve; PA, pulmonary artery; RA, right atrium; RAVV, right atrioventricular valve.

### 3.6 The pulmonary artery, aorta, and coronary arteries

All investigated pulmonary arterial valves had three leaflets of approximately equal size (Figure 6). The leaflets were anchored in 2 dorsal commissures and one ventral commissure (Figure 6j). The aortic valve was approximately of the same size as the pulmonary arterial valve and also exhibited 3 leaflets and commissures (Figure 6). In contrast to the pulmonary arterial valve, the position of the commissures of the aortic valve showed some variation (Figure 6i), such that the dorsal right commissure could be dorsal right (e.g. Ostrich and Chicken, Figure 6a,c), lateral (e.g. Kestrel, Figure 6f), or ventral right (e.g. Budgerigar, Figure 6g). The aorta always gave rise to two coronary arteries, with the right coronary artery originating from the sinus to the right of the ventral commissure and the left coronary artery originating from the sinus to the left of the ventral commissure (Figure 6). Immediately outside the sinus, the right coronary artery gave off a branch that descended into the ventricular septum (Figure 6d) where the flap valve of the right atrioventricular junction merged with the ventricular septum.

### 3.7 The left atrium

In all species, only two pulmonary veins, the left and the right, entered the pericardial cavity (Figure 7). They were of equal size when entering the heart. A sleeve of myocardium was found around both pulmonary veins (Figure 7). With immunohistochemistry, it was confirmed that the myocardium always stopped in the vicinity of the pericardium and the myocardium never extended into the lungs which were located at a much longer distance than the border of the pericardial cavity (Figure 7, Supplementary Figure 6). As the myocardial sleeve stopped, the thickness of the wall of the pulmonary vein decreased around 50% to 75%. The myocardial sleeves around the pulmonary veins, the dorsal ridge and the left atrial shelf formed an antechamber before the body of the left atrium. The prominence of the dorsal ridge was different between species (Table 1). The left atrial shelf formed the ventral boundary of the antechamber and was mostly muscular with some collagen. The free margin of the shelf pointed towards the atrioventricular junction. While the atrial shelf was found in all birds, its extent and thickness as compared to the surrounding wall varied between species (Table 1). The left atrial wall was thin, dominated by a few large pectinate muscles, like the right atrium.

**Figure 7.**
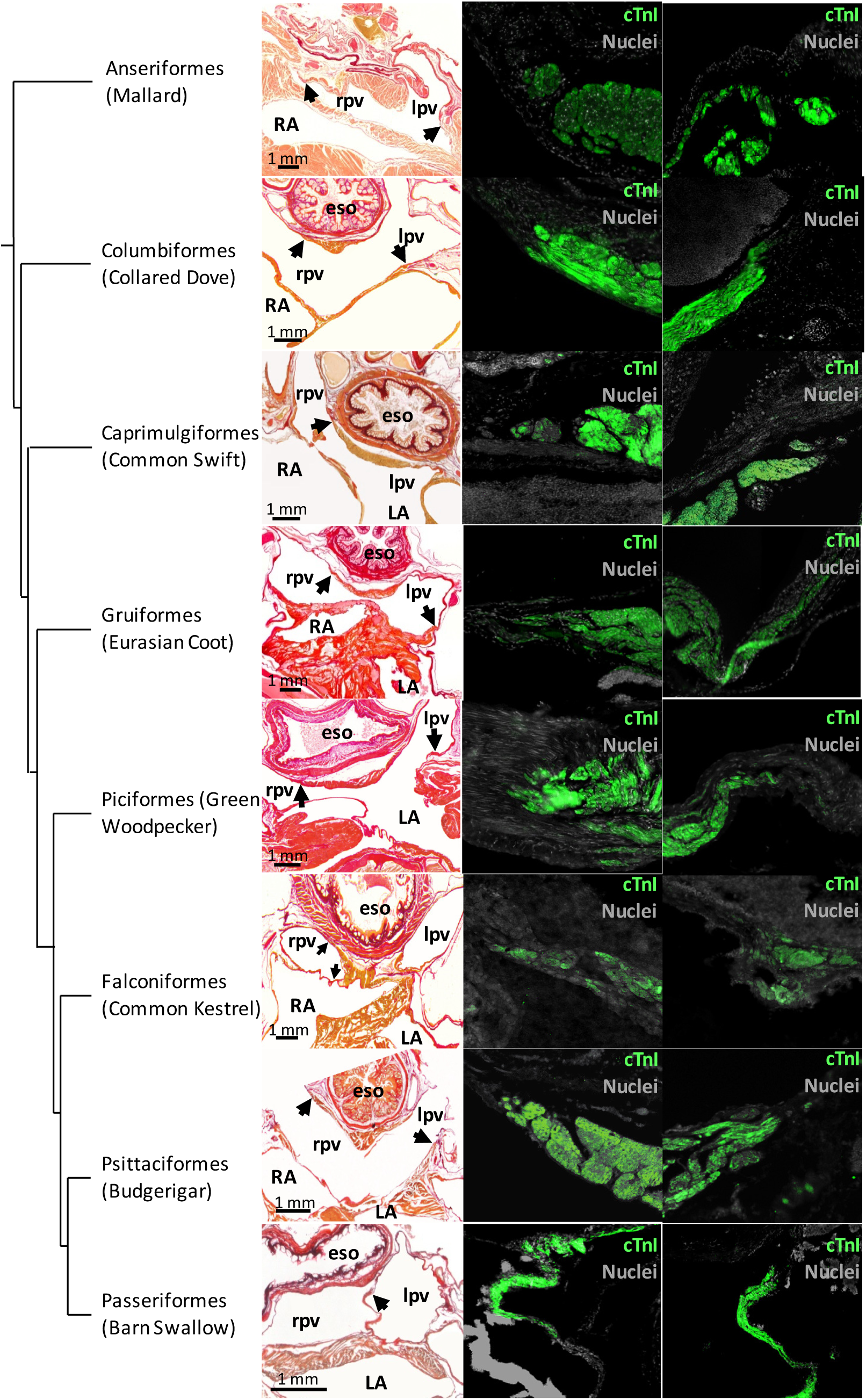
Pulmonary vein myocardium. In the first column picrosirius red stained sections. Black arrows point to the distal-most extent of myocardium. The second and third column show immunohistochemical detection of cTnI at the cites indicated by the arrows. In the Kestrel, only the right pulmonary vein is shown since the left pulmonary vein was damaged during sectioning. Eso, esophagus; LA, left atrium; lpv, left pulmonary vein; RA, right atrium; rpv, right pulmonary vein. In all picrosirius red images, apart from Mallard, the blood has been painted over with white for clarity.

### 3.8 Ventral merger of the atria

In the Budgerigar and the Barn swallow, we found that the left and right atrium embraced the aorta and pulmonary artery such that their walls were merged ventral to the pulmonary artery (Figure 8). The merger spans approximately 1mm from cranial to caudal. In the section series of the Hawfinch heart, in only one section the walls of the left and right atrium merged, which appeared as a band of collagen. In the Lesser redpoll, no merger was found but the two atria were closely juxtaposed. In other orders, the atria were further apart ventral to the pulmonary artery. Only in the Common kestrel the atria were close to each other, but clearly separated by a pad of fatty tissue (Figure 8).

**Figure 8.**
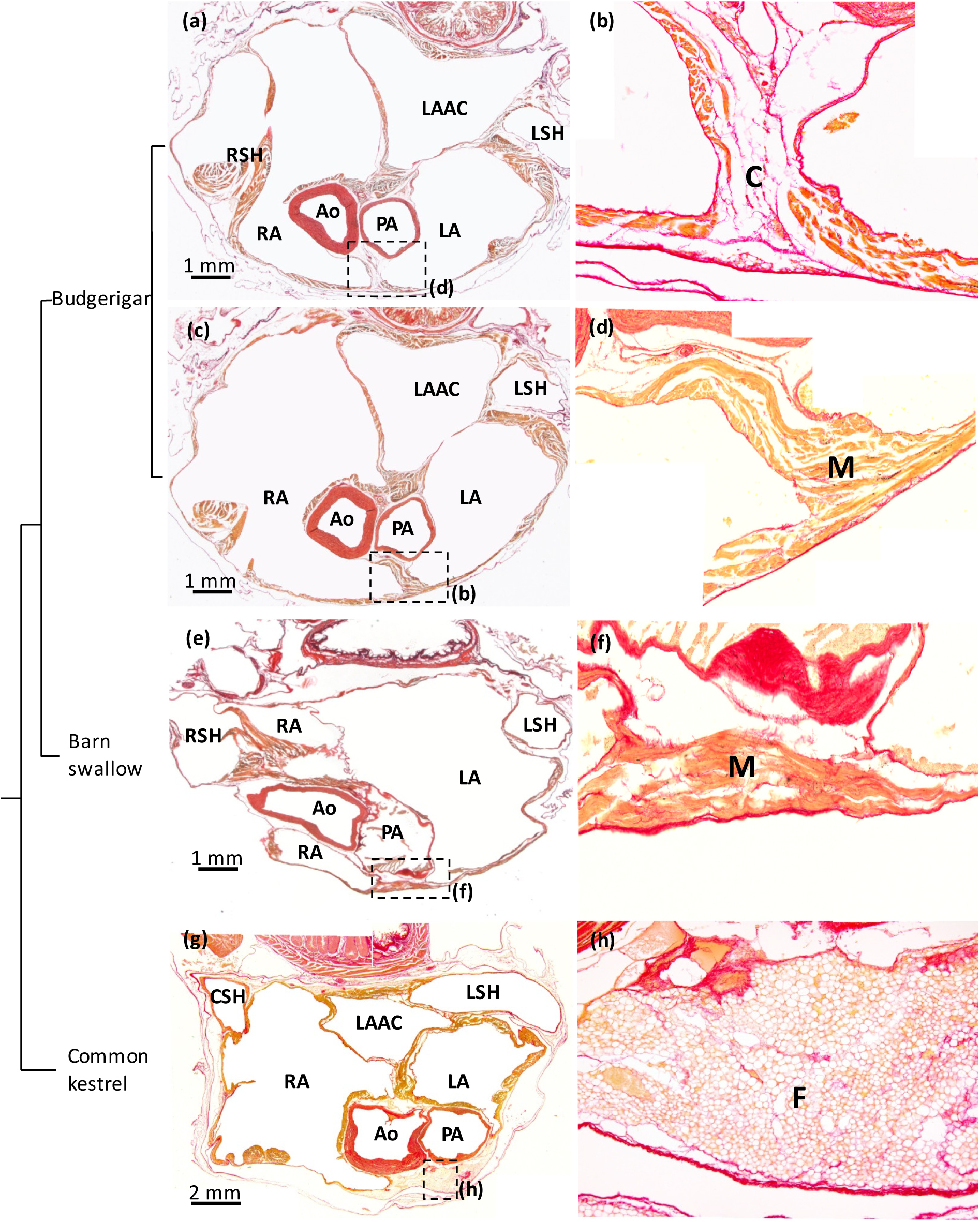
Ventral merger of the atria in budgerigar and barn swallow. (a-d) In the Budgerigar, the atria were merged ventral to the pulmonary artery (PA), by connective tissue (a-b) and myocardium (M) of the left (LA) and right (RA) atrium (c-d). The section in (c) is caudal to (a). In the Barn swallow, the myocardium of the left and right atrium is merged ventral to the pulmonary artery. (g-h) In the Common kestrel, the myocardium of the left and right atrium was not merged ventrally. Ao, aorta; CSH, caudal sinus horn; F, fat; LAAC, left atrial antechamber; LSH, left sinus horn; RSH, right sinus horn. In all images, the blood has been painted over with white for clarity.

## 4 DISCUSSION

In the first subsections below (4.1-4.4), we place our findings in an evolutionary context, leading to the final subsection that argues that the mammalian heart exhibits much more variation than the avian heart (4.5). On this basis, we reject the hypothesis that the transition from ectothermy to endothermy and high cardiac performance is associated with greater variation in cardiac structure.

### 4.1 Gross morphological synapomorphies of the bird heart

Birds are grouped in the archosaur clade, which also includes crocodylians. Hearts of birds share several gross morphological features with crocodylians, and to a lesser extent with other ectothermic reptiles. We show that birds, together with ectothermic reptiles (Jensen et al., 2014b; Cook et al 2017), always have 3 sinus horns (Gasch, 1888; Quiring, 1933), and an atrial septum without a foramen ovale, as previously described (Röse, 1890). As in ectothermic reptiles, in birds the right atrium is more voluminous than the left (Whittow, 1999; Jensen et al., 2014b). In birds and crocodylians, but not in reptiles at large, the left atrioventricular valve comprises membranes anchored in ventricular papillary muscles (Van Mierop, Kutsche, 1985; Lincoln et al., 2004; Cook et al 2017). Birds also share features with crocodylians that, however, are much more developed in birds. This includes the large muscular flap valve in the right atrioventricular junction (Jensen et al., 2014b), the myocardial shelf between the antechamber and left atrial body, which is a meagre flap in crocodiles (Webb, 1979; Cook et al 2017), and the offset between left and right atrioventricular junctions (Cook et al 2017), which we confirm is much more pronounced in birds.

### 4.2. Gross morphological apomorphies of the bird heart

The bird heart is readily distinct from the crocodylian heart, and hearts of other reptiles (Jensen et al., 2014b; Cook et al 2017), by the opening of 2 pulmonary veins with myocardial sleeves to the left atrium. The pulmonary veins empty to a voluminous antechamber with a dorsal ridge of myocardium. A similar antechamber is not seen in crocodylians despite the presence of the muscular shelf (Webb, 1979; Cook et al 2017). In birds, the myocardial shelf is at some distance from the orifices of the pulmonary veins and the left atrioventricular junction. It could prevent regurgitation (Benninghoff, 1933), but given the distances to any orifice, it may guide blood towards the left ventricle instead. The myocardial shelf bears some resemblance to the membrane that divides the left atrium in the human congenital malformation of *cor triatriatum* (Bharucha et al., 2015). Benninghoff (1933) suggested that the myocardial shelf of birds develops from the left pulmonary ridge of the early embryonic atrium. A recent mouse model recapitulates *cor triatriatum* as seen in some patients, but it is not clear whether the setting is the outcome of aberrant development of the left pulmonary ridge (Muggenthaler et al., 2017).

Birds have one aorta with a tricuspid valve, ectothermic reptiles have 2 aortae with bicuspid valves. The atrial walls of birds are dominated by a few, large pectinate muscles coming together in the massive transverse arch in the atrial roof, whereas in ectothermic reptiles the atrial walls consist of a thick meshwork of innumerous tiny trabeculae (Boukens et al^1^; Sedmera et al., 2000). This architecture in birds, parallels the compact organization of their ventricular walls. Functionally, the compact wall architecture likely offers less impedance to the tremendous blood flows that characterize birds and may further allow for higher heart rates by enabling fast electrical activation of the chambers (Boukens et al^1^).

### 4.3 Convergent gross morphological features

The hearts of birds and mammals exhibit convergent features. Since the works of early anatomists, it has been observed repeatedly that birds and mammals have a single aorta, rather than two aortae as in ectothermic reptiles, and that the aortic and pulmonary arterial valve is tricuspid, rather than bicuspid (Benninghoff, 1933; Bartyzel, 2009a; Bartyzel, 2009b). The total number of arterial valve leaflets is thus similar between mammals (6 leaflets), birds (6 leaflets), and ectothermic reptiles (6 leaflets), despite ectothermic reptiles having a left aorta (2 leaflets), a right aorta (2 leaflets), and a pulmonary artery (2 leaflets), reflecting evolutionary conserved morphogenetic processes (Poelmann et al., 2017). Like crocodylians and mammals (Cook et al 2017), there is a full ventricular septum, and the atrioventricular junctions have an offset whereby the left is located cranial to that of the right. Our findings indicate the offset is substantially greater in birds than in crocodylians and mammals. The pronounced offset in birds, may reflect in part that the membranous septum of the ventricle completely myocardializes, whereas the membranous septum of crocodylians exhibits less myocardialization (Jensen et al., 2018) and mammals always have the membranous septum present in the adult heart (Rowlatt, 1990). We confirm the constant presence of a large left atrioventricular junction guarded by membranous leaflets anchored to papillary muscles, and this feature is shared by crocodylians, birds, and mammals (Jensen et al., 2013; Cook et al 2017). The ventricular walls of mammals and birds are much less trabeculated than the ventricles of ectothermic vertebrates (Jensen et al, 2016). Similarly, the atrial walls comprise a thick meshwork of trabeculae in ectotherms, whereas in birds there are some 10 pectinate muscles only as shown here and as previous described in chicken (Sedmera et al., 2000).

### 4.4 Variation between bird hearts

The ventral merger of the atria represents the clearest case for a feature that is not shared by all birds, and it may occur only within passerines and parrots. This feature has not been described previously to the best of our knowledge. In mammals, the left and right atrium can be identified by expression of Pitx2 and Bmp10 respectively (Kahr et al, 2011), but the relative contribution of the left and right atrium to the ventral merger remains to be shown.

Isl1 could not be detected in most specimens, presumably due to the state of tissue preservation, but the Isl1 expressing myocardium had substantial variation in morphology. One extreme was the Mallard, where Isl1 was confined to a nodal structure, not unlike in mammals (Chiodi, Bartolomew, 1967). In contrast, in chicken and the Lesser redpoll, Isl1 was detected in myocardium that, at best, could be distinguished as being slightly thicker than the surrounding walls of the sinus venosus and right atrium. This anatomically poorly defined setting has been recognized previously (Davies, 1930; Chiodi, Bartolomew, 1967; Lamers, De Jong, De Groot, Moorman, 1991) and strongly resembles the pacemaker region of ectothermic reptiles (Jensen et al., 2017). The mammalian sinuatrial node sits at the orifice of the superior vena cava to the right atrium (Keith, Flack, 1907; Davies, 1942; Opthof, 1988; Boyett, Honjo, Kodama 2000; Chandler et al. 2009). Comparative anatomy placed the sinus node of birds at the base of the right leaflet (Davies F, 1930; Chiodi, Bartolomew, 1967), which has been confirmed in chicken by electrophysiological and molecular studies (Moore, 1965; Bressan, Lui, Louie, Mikawa, 2016). Extending those studies, we show in chicken the co-localization of *Bmp2* and Isl1 which is also seen in the pacemaker tissue of Zebrafish (Tessadori et al., 2012) and Anole lizards (Jensen et al., 2017). Note in the Lesser redpoll, however, the Isl1 positive domain extended caudal to the sinuatrial junction (Figure 4j) and not all birds, then, may have pacemaking originating from the base of the right sinuatrial valve leaflet. Also, we confirm the observation by Keith and Flack in the original description of the sinus node, that the sinus node associates with a large coronary artery (Keith, Flack, 1907) (Figure 4b,i).

The state of development of the sinuatrial valve appeared to vary between specimens, and while this was a difficult feature to assess on the basis of histology, the thick valvar margin found in the Green woodpecker was unusual. The extent of myocardium in the pulmonary veins of all specimens was very similar, but the extent of pulmonary venous myocardium may vary in the chicken (Endo et al., 1992). In the Common kestrel, as in other birds of prey (Gasch, 1888), the left leaflet of the sinuatrial valve leaflet was very thin and membranous. In the birds examined with Amira, we found the size of the sinus venosus to be 20% for Mallard and Barn swallow and 30% for Green woodpecker relative to the right atrium. This corresponds to the size relation of the sinus venosus to the atrium in fishes, which is 45% for the White sturgeon and 20% for the Mako shark (Gregory et al., 2004). It appears that the avian sinus venosus has enough muscle to aid atrial filling as in ectotherms, but it is not clear whether it does (Jensen *et al*., 2017). Between specimens, the number of atrial wall pectinate muscles varied. This variation could reflect phylogeny, but size of the heart is likely another factor as we found the hearts of the smallest investigated birds, e.g. barn swallow and lesser redpoll, to have particularly few pectinate muscles. Generally, the position of atrioventricular junctions and arterial bases were fixed, but the aorta did exhibit some rotation in the transverse plane (but much less so than in mammals (Rowlatt, 1990)).

### 4.5 The mammal heart is exceptional varied

Most of the investigated features of the bird heart, exhibit less variation than the same features in mammal hearts. Birds always have 3 sinus horns, whereas mammals may have 2, if the left sinus horn is regressed, or 3. In mammals, the sinuatrial valve can be well-developed and reptile-like as in monotremes, well-developed only around the inferior caval vein and coronary sinus, or almost completely regressed (Rowlatt, 1990; Jensen et al., 2014a). In contrast, in birds both leaflets are always present although their state of development varies as we show here and has been noted before (Benninghoff, 1933).

The atrial septum of monotreme and marsupial mammals develops from the primary septum only. Only in eutherian mammals is the atrial septum formed by the merger of the primary and secondary atrial septum, a process that is revealed in the adult heart by the foramen ovale (Röse, 1890; Runciman, Gannon, Baudinette, 1995; Jensen et al.^2^). In contrast, the atrial septum of birds has no foramen ovale and it develops from the primary septum only (Jensen et al.^2^).

The right atrioventricular junction in monotreme mammals is dominated by a large parietal flap valve, much like in birds and crocodylians, except there is very little if any myocardium in the monotreme valve (Dowd, 1969). In marsupial and eutherian mammals, the right atrioventricular valve is of connective tissue only and typically has 2 or 3 leaflets anchored in papillary muscle (Rowlatt, 1990; Runciman, Baudinette, Gannon, 1992). The right atrioventricular valve in all investigated birds is large, muscular, and without papillary muscle.

In monotreme mammals, the left atrium receives a single pulmonary vein, like in ectothermic reptiles, whereas between genera of marsupial and eutherian mammals, the number of pulmonary veins can vary between 1 and 7 and myocardial sleeves may be absent, short or extensive (Rowlatt, 1990). In contrast, 2 pulmonary veins connecting to the left atrium is a “konstant” feature of birds (Benninghoff, 1933). We confirm Benninghoff’s observation and further show that the pulmonary veins have myocardial sleeves that extend to the pericardial boundary. In mammals, the myocardial sleeves are often the cause of atrial fibrillation through automaticity and reentry (Haïssaguerre et al., 1988; Chen et al., 1999; Tsuneoka, Koboyashi, Honda, Namekata, Tanaka, 2012), and it remains to be shown whether the pulmonary veins of birds are similarly arrhythmogenic.

In mammals, there is always a ventricular membranous septum which varies in size between genera and may be covered by a layer of myocardium (Rowlatt, 1990). Birds do form a membranous septum in development, but it undergoes myocardialization and is subsequently lost. It may be growth of this myocardium of mesenchymal origin that contributes to the greater offset between left and right atrioventricular junctions.

The relative position of the atrioventricular junctions and the base of the pulmonary artery and the aorta is not fixed across mammals (Rowlatt, 1990). The left and right atrioventricular junctions may be juxtaposed as in tree squirrels or far apart as in pygmy sperm whales. This, in turn, appears to reflect the process of aortic wedging, the extent to which the aorta has moved from its embryonic position on the right and towards the left ventricle (Cook et al 2017). In contrast, we found in all birds the same relative position of the atrioventricular junctions and the base of the pulmonary artery and the aorta. Only the commissures of the aortic valve appeared to exhibit some relative rotation, but such rotation appears much greater in mammals (Rowlatt, 1990).

The bird heart has several features that set it apart from that of ectothermic reptiles but these features do not exhibit much variation. In contrast, the features that set mammals apart from their fellow amniotes, exhibit substantial variation. Therefore, our data do not support the hypothesis that the transition from ectothermy to endothermy associated with greater variation in cardiac design. While its clear there is a common building plan to the amniote heart (Keith, Flack, 1907; Olson, 2006; Jensen, Wang, Christoffels, Moorman, 2013), the basis of the morphogenetic plasticity of mammals remains to be explored.

## 5 CONCLUSION

We assessed 15 features of and around the atria of hearts from 13 orders of birds. Most features were surprisingly constant in appearance between orders, even those that were not shared with ectothermic reptiles and thus considered apomorphic. This suggests, in contrast to what was hypothesized, that the transition from ectothermy to endothermy and the concomitant evolution of a high-performance heart, does not necessarily lead to the extraordinary degree of variability in cardiac design that is observed in mammals.

## ACKNOWLEDGMENTS

Dr. J.C.M. Schouten was an excellent help in procuring hearts for the study.

## AUTHOR CONTRIBUTIONS

ORCID

*Bjarke Jensen* https://orcid.org/0000-0002-7750-8035

## REFERENCES

Bartyzel B.J., (2009a). Morphology of the pulmonary valve (valva trunci pulmonali) in chosen species of domestic and wild birds using imaging methods. Bulletin of the Veterinary Institute in Pulawy 53: 303–308

Bartyzel B. J., (2009b). The aortic valve and other heart structures of selected species of sea birds in a morphological and imaging scope. Electronic journal of polish agricultural universities 12.4

Benninghoff, A., (1933). Das Herz. In L. Bolk, E. Goppert, E. Kallius and W. Luhosch (Eds), Handbuch der Vergleichenden Anatomie der Wirbeltiere, 6: 467–556. Berlin and Wien: Urban und Schwarzenberg.

Bharucha T., Spicer D. E., Mohun T. J., Black D., Henry G. W., Anderson R. H., (2015). Cor triatriatum or divided atriums: which approach provides the better understanding? Cardiology in the Young, 25(2):193–207

Boyett M. R., Honjo H., Kodama I., (2000). The sinoatrial node, a heterogeneous pacemaker structure. Cardiovascular Research, 47(4): 658–687.

Bressan M., Lui G., Louie J. D., Mikawa T., (2016) Cardiac pacemaker development from a tertiary heart field. In: Nakanishi T., Markwald R., Baldwin H., Keller B., Srivastava D., Yamagishi H. (eds) Etiology and Morphogenesis of Congenital Heart Disease. Springer, Tokyo. pp 281–288

Calkins H., Brugada J., Packer D. L., Cappato R., Chen S.-A., Crijns H. J. G., Damiano Jr. R. J., (…), Shemin R. J., (2007). HRS/EHRA/ECAS Expert Consensus Statement on Catheter and Surgical Ablation of Atrial Fibrillation: Recommendations for Personnel, Policy, Procedures and Follow-Up. Heart Rhythm, 4(6): 816–861.

Carmona, R., Ariza, L., Cañete, A., Muñoz-Chápuli, R., (2018). Comparative developmental biology of the cardiac inflow tract. Journal of Molecular and Cellular Cardiology, 116, 155–164

Chandler N. J., Greener I. D., Tellez J. O., Inada S., Musa H., Molenaar P., Difrancesco D., Baruscotti M., Longhi R., Anderson R. H., Billeter R., Sharma V., Sigg D. C., Boyett M. R., Dobrzynski H., (2009). Molecular architecture of the human sinus node: insights into the function of the cardiac pacemaker. Circulation, 119(12): 1562–1575.

Chen S.A., Hsieh M. H., Tai C. T., Tsai C. F., Prakash V. S., Yu W. C., Hsu T. L., Ding Y. A., Chang M. S., (1999) Initiation of atrial fibrillation by ectopic beats originating from the pulmonary veins: Electrophysiological characteristics, pharmacological responses, and effects of radiofrequency ablation. Circulation, 100(18):1879–1886.

Chiodi, V., Bortolami, R., (1967) The conducting system of the vertebrate heart. Calderine, Bologna.

Cook A. C., Tran V. H., Spicer D. E., Rob J. M. H., Sridgaran S., Taylor A., Anderson R. H., Jensen B., (2017) Sequential segmental analysis of the crocodilian heart. Journal of Anatomy, 231(4): 484–499.

Crossley G. H., Sorrentino R. A., Exner D. V., Merliss A. D., Tobias S. M., Martin D. O., Augostini R., Piccini J. P., Schaerf R., Li S., Miller C. T., Adler S. W., (2016) Extraction of chronically implanted coronary sinus leads active fixation vs passive fixation leads. Heart Rhythm, 13(6):1253–1259. doi: 10.1016/j.hrthm.2016.01.031.

Davies, F., (1930). The Conducting System of the Bird’s Heart. Journal of Anatomy, 64(Pt 2), 129–146.7.

Davies, F., (1942). The conducting system of the vertebrate heart. British Heart Journal 4, 66–76

Dowd, D. A., (1969). The coronary vessels and conducting system in the heart of monotremes. Cells Tissues Organs, 74.4: 547–573

Endo H., Kurohmaru, M., Nishida, T., Hayashi, Y. (1992) Cardiac musculature of the cranial and caudal venae cavae and the pulmonary vein in the fowl. Journal of Veterinary Medical Science, 54, 479–484

Gasch F. R., (1888) Beiträge zur vergleichenden Anatomie des Herzens der Vögel und Reptilien. Archiv für Naturgeschichte, 54(1):119–154.

Green R.E., Braun E.L., Armstrong J., Earl., Nguyen N., Hickey G., … Ray D.A., (2014). Three crocodilian genomes reveal ancestral patterns of evolution among archosaurs. Science, 346.6215.

Gregory J. A., Graham J. B., Cech J. J. Jr., Dalton N., Michaels J., Chin Lai N., (2004). Pericardial and pericardioperitoneal canal relationships to cardiac function in the white sturgeon (Acipenser transmontanus). Comparative Biochemistry and Physiology Part A: Molecular & Integrative Physiology, 138(2): 203–213.

Haïssaguerre M., Jaïs P., Shah D. C., Takahashi A., Hocini M., Quiniou G., Garrigue S., Mouroux A., Métayer P., Clémenty J., (1998). Spontaneous initiation of atrial fibrillation by extopic beats originating in the pulmonary veins. The New England Journal of Medicine, 339(10):659–666.

Hamburger V., Hamilton H.L., (1951) A series of normal stages in the development of the chick embryo. Journal of Morphology 88:1 49–92.

Jensen B., Agger P., de Boer B. A., Oostra R. J., Pedersen M., van der Wal A. C., Planken N. R., Moorman A. F., (2016). The hypertrabeculated (noncompacted) left ventricle is different from the ventricle of embryos and ectothermic vertebrates. Biochimica et Biophysica Acta Molecular Cell Research, 1863 (7 Pt B): 1696–1706 https://doi.org/10.1016/j.bbamcr.2015.10.018.

Jensen B., Boukens B. J. D., Wang T., Moorman A. F. M., Christoffels V. M., (2014). Evolution of the sinus venosus from fish to human. Journal of Cardiovascular Development and Disease, 1:14–28.(a)

Jensen B., Boukens B. J. D., Crossley D. A., Conner J., Mohan R. A., van Duijvenboden K., Postma A. V., Gloschat C. R., Elsey R. M., Sedmera D, Efimov I. R., Christoffels V. M., (2018). Specialized impulse conduction pathway in the alligator heart. eLife 7: e32120

Jensen B., Moorman A. F. M., Wang T., (2014). Structure and function of the hearts of lizards and snakes. Biological Reviews of the Cambridge Philosophical Society, 89(2):302–336.(b)

Jensen B., van den Berg G., van den Doel R., Oostra R. J., Wang T., Moorman A. F. M., (2013). Development of the Hearts of Lizards and Snakes and Perspectives to Cardiac Evolution. PLoS One, 8(6): e63651.

Jensen B., Vesterskov S., Boukens B. J. D., Nielsen J. M., Moorman A. M. F., Christoffels V. M, Wang T., (2017). Morpho-functional characterization of the systemic venous pole of the reptile heart. Scientific reports, 7 doi: 10.1038/s41598-017-06291-z.

Jensen B., Wang T., Christoffels V. M., Moorman A. F. M., (2013). Evolution and development of the building plan of the vertebrate heart. Biochimica et Biophysica Acta Molecular Cell Research, 1833(4): 783–794.

Kahr P. C., Piccini I., Fabritz L., Greber B., Schöler H., Scheld H. H., Hoffmeier A., Brown N. A., Kirchhof P., (2011). Systematic analysis of gene expression differences between left and right atria in different mouse strains and in human atrial tissue. PLoS One, 6(10).

Keith A., Flack M., (1907), The form and nature of the muscular connections between the primary divisions of the vertebrate heart. Journal of Anatomy and Physiology, 41(Pt 3), 172.

Lamers, W. H., De Jong F., De Groot I. J., Moorman A. F., (1991). The development of the avian conduction system, a review. European journal of morphology 29.4: 233–253.

Lincoln J., Alfieri C. M., Yutzey K. E., (2004) Development of heart valve leaflets and supporting apparatus in chicken and mouse embryos. Developmental Dynamics 230.2: 239–250

Mansour M., Holmvang G., Sosnovik D., Migrino R, Abbara S., Ruskin J., Keane D., (2004). Assessment of pulmonary vein anatomic variability by magnetic resonance imaging: implications for catheter ablation techniques for atrial fibrillation. Journal of Cardiovascular Electrophysiology, 15(4): 387–393

Mommersteeg M. T. M., Brown N. A., Prall O. W. J., de Gier-de Vries C., Harvey R. P., Moorman A. F. M., Christoffels V. M., (2007). Pitx2c and Nkx2-5 are required for the formation and identity of the pulmonary myocardium. Circulation Research, 101(9): 902–909.

Moore, E. N. (1965). Experimental electrophysiological studies on avian hearts. Annals of the New York Academy of Sciences, 127(1), 127–144

Muggenthaler M. M. A., Chowdhury B., Hasan S. N., Cross H. E., Mark B., Harlalka G. V., Patton M. A., (…), Chioza B. A., (2017). Mutations in HYAL2, Encoding Hyaluronidase 2, Cause a Syndrome of Orofacial Clefting and Cor Triatriatum Sinister in Humans and Mice. PLoS Genetics, 13(1) art. no. e1006470.

Nathan H., Eliakim M., (1966). The junction between the left atrium and the pulmonary veins. An anatomic study of human hearts. Circulation, 34(3): 412–422.

Olson, E. N., (2006). Gene regulatory networks in the evolution and development of the heart. Science 313.5795: 1922–1927

Opthof, T., (1988). The mammalian sinoatrial node. Cardiovascular drugs and therapy 1.6: 573–597

Poelmann, R. E., Gittenberger-de Groot, A. C., Biermans, M. W. M., Dolfing, A. I., Jagessar, A., van Hattum, S., … Richardson, M. K., (2017). Outflow tract septation and the aortic arch system in reptiles: lessons for understanding the mammalian heart. EvoDevo, 8, 9. http://doi.org/10.1186/s13227-017-0072-z

Poelmann, R. E., Mikawa T., Gittenberger-De Groot, A. C., (1998). Neural crest cells in outflow tract septation of the embryonic chicken heart: differentiation and apoptosis. Developmental Dynamics 212.3: 373–384

Quiring D. P., (1933). The development of the sino-atrial region of the chick heart. Journal of Morphology, 55(1):81–118.

Röse C., (1890). Beitrage zur vergleichenden Anatomie des Herzens der Wirbeltiere. Gegenbaurs Morphologisches Jahrbuch, 16: 27–96

Rowlatt, U., (1968). Functional Morphology of the Heart in Mammals. Integrative and Comparative Biology, 8;2(1):221–229.

Rowlatt, U., (1990). Comparative anatomy of the heart of mammals. Zoological Journal of the Linnean Society, 1;98(1):73–110.

Runciman S. I. C., Baudinette R. V., Gannon B. J., (1992). The Anatomy of the Adult Marsupial Heart - an Historical Review. Australian Journal of Zoology, 40:21–34

Runciman S. I. C., Gannon B. J., Baudinette R. V., (1995). Central cardiovascular shunts in the perinatal marsupial. Anatomical Record, 243:71–83

Sedmera D., Pexieder T., Vuillemin M., Thompson R.P., Anderson R.H., (2000) Developmental patterning of the myocardium. Anatomical Record, 1; 258(4): 319–337.

Somi S., Buffing A. A. M., Moorman A. F. M., van den Hoff M. J. B., (2004). Dynamic patterns of expression of BMP isoforms 2, 4, 5, 6, and 7 during chicken heart development. The Anatomical Record 279.1, 636–651.

Tattersall G. J., (2016). Reptile thermogenesis and the origins of endothermy. Zoology 119.5: 403–405

Tessadori F., van Weerd J. H., Burkhard S. B., Verkerk A.O., de Pater E., Boukens B. J., Vink A., Christoffels V. M., Bakkers J., (2012). Identification and Functional Characterization of Cardiac Pacemaker Cells in Zebrafish. PLoS One, 7(10).

Tsuneoka Y., Kobayashi Y., Honda Y., Namekata I., Tanaka H., (2012). Electrical activity of the mouse pulmonary vein myocardium. Journal Pharmacological Science, 119:287–292.

van den Berg G., Abu-Issa R., de Boer B. A., Hutson M. R., de Boer P. A., Soufan A. T., Ruijter J. M., Kirby M. L., van den Hoff M. J., Moorman A. F., (2009). A caudal proliferating growth center contributes to both poles of the forming heart tube. Circulation research 104.2: 179–188.

van den Hoff M. J. B., Kruithof B. P. T., Moorman A. F. M., Markwald R. R, Wessels A., (2001). Formation of myocardium after the initial development of the linear heart tube. Developmental biology, 240.1, 61–76.

Van Mierop L. H. S., (1967). Location of pacemaker in chick embryo heart at the time of initiation of heartbeat. American Journal of Physiology-Legacy Content 212.2: 407–415

Van Mierop L. H. S., Kutsche L. M., (1985). Development of the ventricular septum of the heart. Heart and Vessels, 1(2): 114–119. https://doi.org/10.1007/BF02066358

Warren, W. C., Hillier, L. W., Marshall Graves, J. A., Birney, E., Ponting, C. P., Grützner, F., … Wilson, R. K., (2008). Genome analysis of the platypus reveals unique signatures of evolution. Nature, 453(7192), 175–183.

Webb, G., (1979). Comparative cardiac anatomy of the Reptilia. III. The heart of crocodilians and an hypothesis on the completion of the interventricular septum of crocodilians and birds. Journal of Morphology, 161(2): 221–240.

Whittow G. C., (1999). Sturkie’s Avian Physiology (Fifth Edition). Academic Press; Cambridge, Massachusetts.

Wyneken J., (2009). Normal reptile heart morphology and function. Veterinary Clinics of North America: Exotic Animal Practice, 12(1): 51–63.

Boukens et al. in press^1^ and Jensen et al., in press, Anat Rec^2^ are unpublished at this time and therefore not full references.

